# Estimating Organism Abundance Using Within-Sample Haplotype Frequencies of eDNA Metabarcoding Data

**DOI:** 10.1101/2025.06.30.662414

**Authors:** Pedro FP Brandão-Dias, Gledis Guri, Megan Shaffer, Elizabeth Andruszkiewicz Allan, Ryan P Kelly

## Abstract

Environmental DNA (eDNA) metabarcoding provides powerful insights into species presence and community composition, but remains limited in its ability to quantify species abundance or structure. Here, we show that deviation between observed haplotype frequencies within a given sample and the population haplotype frequencies can be used to infer the number of individual contributors to an eDNA sample. We also lay out the theory for how population haplotype frequencies can be approximated from eDNA data alone, enabling broad applicability even in the absence of tissue-based references. We then present an estimator to derive the number of individual contributors to a given eDNA sample and validate its performance using simulations with variable allele frequencies and noise. Our framework demonstrates that differences between expected and observed frequencies carry meaningful biological information in eDNA data. Our results show that the number of contributors can be recovered under a range of conditions, particularly with hypervariable markers and sufficient sampling. This approach complements existing molecular methods and opens a new avenue for inferring abundance from eDNA metabarcoding datasets.

## INTRODUCTION

Most vertebrate environmental DNA (eDNA) studies have generally focused on either single-species PCR, which enables precise detection and quantification of target taxa eDNA, or on metabarcoding, which provides broader snapshots of community composition but typically yields only semi-quantitative data. Although both approaches have greatly advanced our understanding of biodiversity, the limited quantitative information available from eDNA metabarcoding data hampers our ability to accurately monitor species abundances and dynamics at larger scales. Thus, obtaining more robust quantitative information from metabarcoding data could combine the major advantages of both methods, combining precise quantification with the ability to detect and monitor broad sections of biodiversity, even without prior knowledge.

While read counts in metabarcoding data are often interpreted quantitatively in the literature, and superficial correlation between sequencing read number and species abundance may exist in metabarcoding data (Elbrecht & Leese, 2015; Lamb et al., 2022), even under identical sequencing depths, the number of reads assigned to a particular organism is affected by many biases (Gold et al., 2023; Shaffer et al., 2025), which may lead to incorrect interpretations. A key source of bias is amplification bias, which is due to how well a given organism’s DNA is amplified in PCR, often as a result to sequence mismatches with primers (Shaffer et al., 2025; Sipos et al., 2007). This can significantly alter the representation of species proportions in metabarcoding results (Elbrecht & Leese, 2015; Polz & Cavanaugh, 1998; Shelton et al., 2023). Additionally, because metabarcoding data is compositional, an increase in the abundance of other organisms within the sample will reduce the read depth and proportion of reads for others in the sample, even if their absolute DNA quantities remain unchanged (Shelton et al., 2023). Thus, although semiquantitative analyses of metabarcoding data often provide valuable insights (Guri, Westgaard, et al., 2024) metabarcoding data should not be considered as strictly quantitative without prior corrections (Elbrecht & Leese, 2015; Shelton et al., 2023).

Recent statistical advances have enabled quantitative interpretations of metabarcoding data, allowing for more accurate and simultaneous assessments of DNA quantities of multiple organisms in environmental samples (Guri, Shelton, et al., 2024). By pairing metabarcoding with amplification efficiency data from mock communities— controlled mixtures of known DNA sequences—and quantitative PCR (qPCR) data for a single frequently observed species, we can correct for the amplification biases and then derive absolute DNA quantifications from compositional metabarcoding data (Allan et al., 2023; Shelton et al., 2023). However, even if we obtain precise information on the quantity of eDNA within samples, the relationship between total eDNA and number of organisms is a complex function of organism size, metabolism, biotic and abiotic variables of the system, spatiotemporal dynamics of eDNA decay and transport, among other sources of variance (Yates et al., 2025), all of which complicate absolute abundance estimations across diverse organisms. Therefore, obtaining absolute quantities of DNA within environmental samples, be that by quantitative PCR or correcting metabarcoding data, is still several steps away from determining organism abundance or biomass in a region.

However, metabarcoding data offers more information beyond species composition and read counts, which has yet to be fully explored - the presence of multiple amplicon sequence variants (ASVs) within a detected species. Water samples collected for eDNA analyses represent a mix of contributions of DNA from multiple individuals, thus being a natural sample of genetic diversity within an area. How much DNA each individual organism contributes to an eDNA sample will depend on several factors including their size (Maruyama et al., 2014; Yates et al., 2020) and the distance in space and time between target organism and sampling location (Harrison et al., 2019; Laporte et al., 2020). Nonetheless, when individuals possess different alleles at a given locus, the eDNA sample will contain a mixture of these alleles, with their relative frequencies proportional to the contributions of each individual. Importantly, these allele frequencies are not random, but reflect the aggregated contributions of all individuals shedding DNA into the sample. Consequently, as the number of contributors increases, the number of observed alleles increases (Ai et al., 2025; Andres et al., 2021), and, importantly, the observed allele frequencies within a sample are expected to converge toward the population frequencies of these alleles (Figure 1, Figure S1).

**Figure 1:**
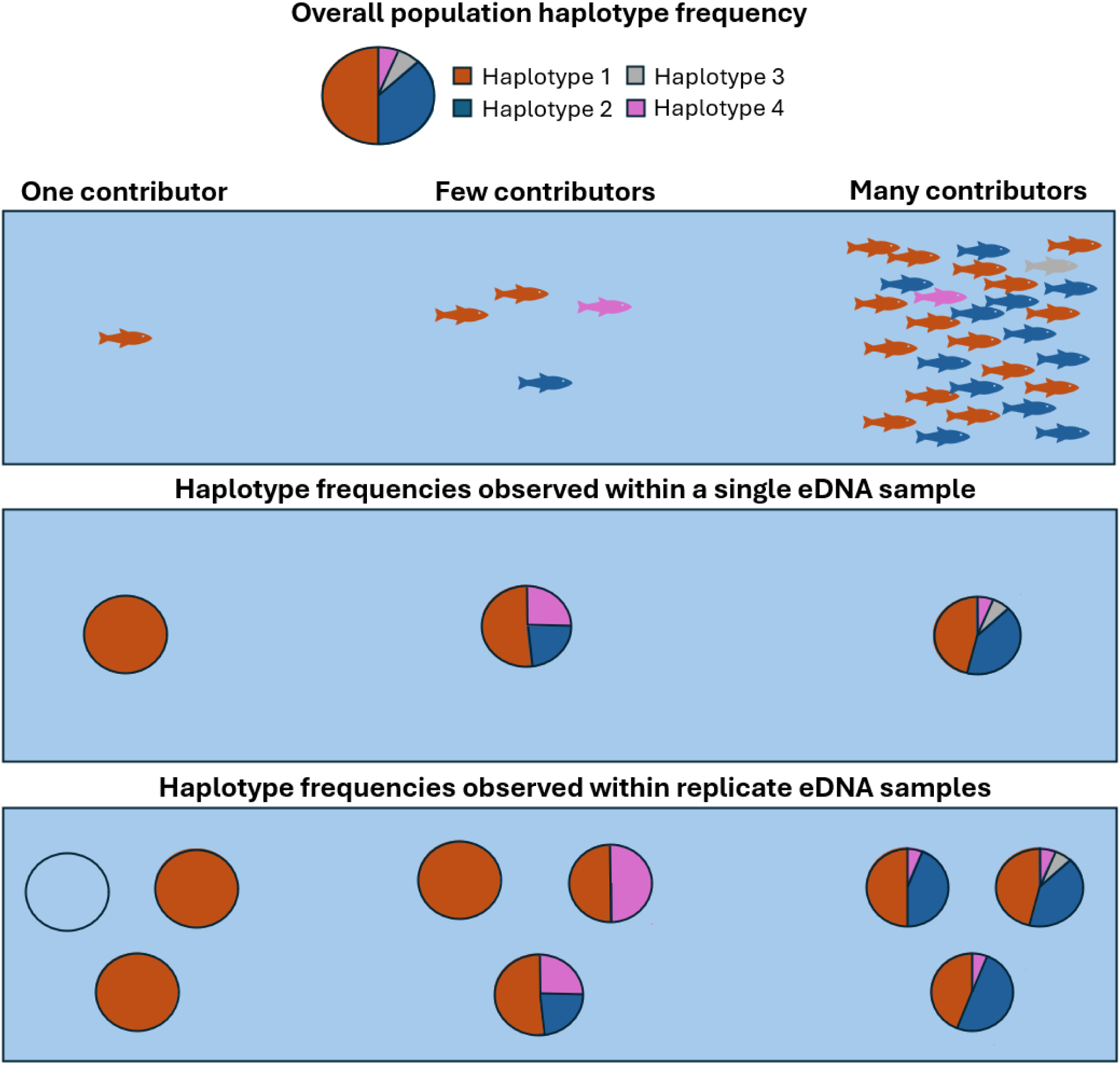
Convergence of observed eDNA haplotype frequencies toward population frequencies with increased number of contributors. The pie chart on the top left depicts the population haplotype frequencies, corresponding to the distribution of haplotypes (Haplotypes 1–4) represented by colors. With fewer contributors, the observed haplotype frequencies are quite different from the population frequencies, but with more contributors, eDNA from a greater diversity of individuals is captured, and observed haplotype frequencies increasingly align with population frequencies. With a single eDNA sample, the difference between within sample haplotype frequencies and the population frequencies can be used to infer the number of contributors. When multiple samples are available, the difference between samples will also offer an additional level of information regarding the variability of the observation process.

Under panmixia, the haplotype composition of an eDNA sample reflects a multinomial draw from the underlying population haplotype frequencies. Each individual contributes DNA carrying specific haplotypes, and the sample captures a subset of this diversity, much like drawing colored marbles from a jar. As the number of contributors increases, sampling variance naturally decreases, and observed haplotype frequencies converge toward population-level values. When few individuals contribute, frequencies deviate more strongly due to stochastic sampling (Figs 1, S1). These deviations are not just noise—they carry information. The extent to which observed haplotype frequencies differ from population expectations can reveal how many individuals contributed: large deviations suggest few contributors; close matches point to many.

Similar but distinct principles have been previously applied to eDNA samples (Ai et al., 2025; Andres et al., 2021; Yoshitake et al., 2019). The core realization behind all of these methods is that an eDNA sample is a complex, mixed genetic signal from multiple contributors, but these methods rely on different subsets of the information encoded in the haplotypes detected. Conceptually, a mixed eDNA sample provides three tiers of information: (1) the number of distinct haplotypes present, (2) the identity of those haplotypes, and (3) their observed frequencies within the sample.

The first tier relies only on haplotype count - how many distinct haplotypes are detected - without considering their identities or frequencies. Yoshitake et al (2019) demonstrated that the number of haplotypes from a hyper-variable region can be used to infer number of individuals in an eDNA sample. More recently, Ai et al. (2025) demonstrated that, as expected, number of haplotypes correlates strongly with the number of individual contributors, making a frequency-agnostic estimator of contributor abundance. While this method is elegant in its simplicity, its informativeness is dependent on the underlying but unknown haplotype frequency distribution. For instance, consider a population with three alleles (a, b, c) at frequencies 0.87, 0.12, and 0.01. A sample containing only allele a or only allele c would produce identical abundance estimates under this method, despite their vastly different probabilities of arising from multiple contributors. While useful in some contexts, this approach cannot generalize well across markers and populations with varying allele frequencies.

The second tier of information, the identity of haplotypes, was previously explored to determine number of contributors to eDNA samples by Andres et al. (Andres et al., 2021; Andres, Lodge, & Andrés, 2023; Andres, Lodge, Sethi, et al., 2023). Their method builds on earlier work with mixed genetic samples from forensic settings (Haned et al., 2011; Sethi et al., 2019; Weir et al., 1997), but it does not consider the frequency of haplotypes within samples, only their identity and respective known population haplotype frequencies. Therefore, because it discards within-sample frequency information, it assumes those are irrelevant – a fair assumption on a forensic scenario where the method originates (Paoletti et al., 2005; Weir et al., 1997), but incompatible with eDNA ecology. While this model may be particularly useful when frequency data are extremely noisy, such as under strong stochastic effects or technical error (e.g., the cumulative noise shown in Fig. 2f), if allele frequencies are too noisy to be informative, then treating presence/absence as more reliable is equally problematic, especially for rare alleles. Thus, there is little justification for ignoring frequency information while still considering haplotype identity. A key methodological advance introduced here is the use of the third tier of information in the within-sample haplotype frequencies.

**Figure 2:**
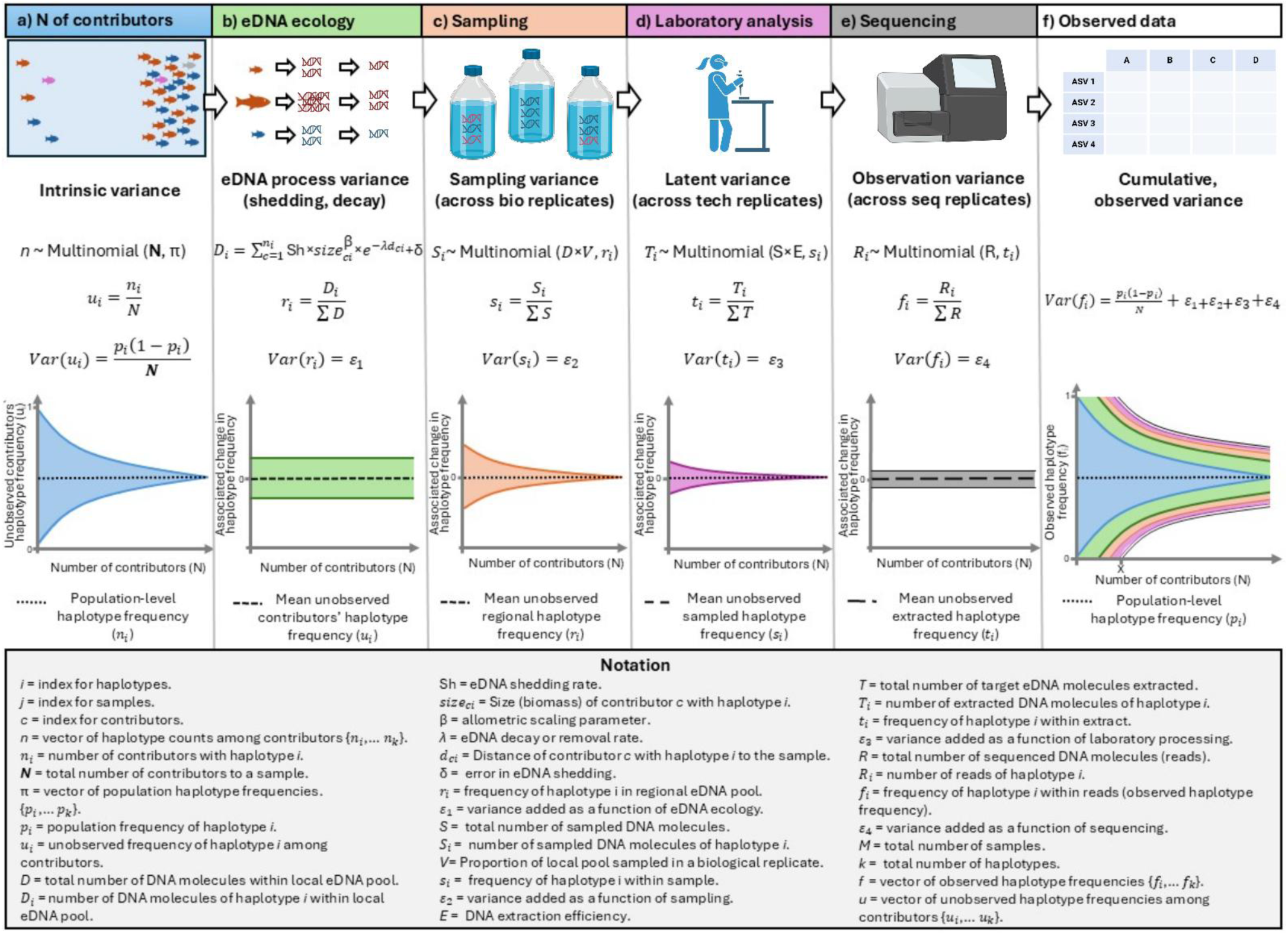
Propagation of variance in haplotype frequencies observed within an environmental DNA sample. Panel (a) illustrates the inherent process of eDNA production and distribution in the natural environment, where the variability in haplotype frequencies is intrinsically higher in regions with fewer contributors (*N*). This baseline variance allows one to infer the original number of contributors by examining haplotype frequency variability across samples. In panel (b) we observe the eDNA process variance, where different biomass and other parameters result in differential shedding rate, or distance between contributor and sample results in eDNA decaying or getting transported away before getting samples. The result is that different contributors provide a variable amount of eDNA to the regional DNA pool D, affecting regional haplotype frequencies (*ri*). Panel (c), *Di* are sampled, yielding intermediate, unobserved sampled frequencies for each halotype (*si*). Panel (d) depicts the laboratory processes—such as DNA extraction, pipetting—that further add noise, resulting in intermediate extracted haplotype frequencies (*ti*). In panel (e), the sequencing step is represented as a multinomial sampling process, where the variance of the observed haplotype frequencies (*fi*) is a function of the intermediate frequencies (*ti*) and is inversely proportional to the sequencing read depth (R); deeper sequencing results in lower variance at this step. Finally, panel (f) shows that the final observed haplotype frequencies incorporate the cumulative variance from all stages—from natural eDNA distribution through sampling, laboratory processing, and sequencing. Point X on the x-axis shows the N at which variance becomes indistinguishable due to added noise.

A fundamental requirement of many of the methods that use haplotypes or alleles to estimate the number of contributors, in additional to many populations genetic methods, is knowledge of the population’s haplotype or allele frequencies (π), a level of detail rarely available for organisms detected through eDNA metabarcoding and often cited as a major barrier to applying these methods. In practice, this creates a conceptual paradox: if a population has been extensively sampled to estimate haplotype frequencies, then its abundance is likely already known, making further eDNA-based inference redundant.

However, numerous independent studies over the years have demonstrated that population frequencies can be potentially inferred directly from eDNA data. For example, with a rare dataset, Sigsgaard et al. (2016) showed that haplotypes recovered from multiple whale shark eDNA detections matched frequencies observed in tissue samples. Uchii et al. (2016) found that carp haplotype frequencies from environmental samples aligned with those in controlled systems. Andres, Lodge, & Andrés, (2023) also recovered accurate nuclear and mitochondrial alleles for gobies, though eDNA also sampled rare haplotypes that were absent in tissue frequencies. Wakimura et al. (2023) showed that haplotype frequencies of an endangered fish species recovered from eDNA matched those of traditional sampling after proper filtering. In the following study, Wakimura et al. (2025) found again that eDNA recovered mtDNA haplotypes of an invasive fish matched the tissue-derived frequencies from previous studies. Finally, Parsons et al. (2025) found no bias in the frequency of haplotypes of porpoises generated from eDNA samples when compared to tissue samples.

Beyond eDNA’s repeated success in recovering population haplotypes in empirical studies where tissue-derived frequencies were available, there are strong theoretical reasons to expect that eDNA samples capture a representative snapshot of a population’s genetic diversity. As discussed above, eDNA in any environmental sample is inherently a mixture of DNA from multiple individual contributors. When samples are taken deliberately near a particular individual, such as in Parsons et al. (2025) who sample fluke prints of marine mammals, the haplotypes recovered may reflect that nearby individual’s genetic material, meaning the presence of haplotypes is informative, but their frequencies are not. However, when samples are evenly dispersed across space, capturing DNA from many individuals, and samples encompass the same population, both the presence and relative frequencies of haplotypes can provide meaningful information. In such cases, each sample reflects an indirect, uneven, and stochastic draw from the population’s genetic pool, shaped by local density, movement, and shedding dynamics. Hence, the average frequency of haplotypes across all samples should be a fair approximation of the population frequencies, provided enough samples are drawn and no haplotype-specific observation bias.

In light of these theoretical expectations, we present an approach to estimate the population frequency of haplotypes with eDNA samples, and subsequently estimate the number of contributors to each eDNA sample. These methods rely exclusively on the observed within-sample frequencies, expanding the utility of metabarcoding data beyond presence-absence assessments to provide estimates of organismal abundance in mixed samples when multiple ASVs are identified for a target organism. In this study, we describe the methods in detail, and validate their potential and limitations using simulated eDNA data. While not a direct measure of absolute abundance, this framework complements other quantitative methods by offering additional insights into the interpretation of eDNA metabarcoding results.

## METHODS

As described above, the variability of haplotype frequencies within eDNA observations relative to population frequencies carry information about the number of contributors (*N*) to each sample. We extract this information in two steps. First, we use the full, spatially replicated eDNA data set to derive a population-level haplotype frequency vector π: each sample contributes equally to a leave-one-out mean (see below). Second, we offer a normal approximation maximum-likelihood estimator (MLE) to obtain both a point estimate and confidence interval for *N*. The sections below will walk through these derivations.

### Common Mathematical Framework

Let ***π*** = *(pi,…,pK)* denote the population frequencies of haplotypes *i* ∈ {1, … *k*} at the same locus, such that all the frequencies sum to 1. We assume that independent eDNA samples *j* have been collected and denote by *N_j_* the total (unknown) number of individual contributors to sample *j*. Under panmixia, the set of individual contributors for each sample will follow a multinomial draw:

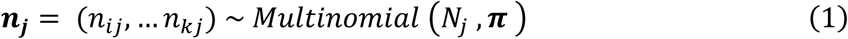

where *n_ij_* is the unobserved number of contributors in sample *j* that carry haplotype *i*. Thus, the relative frequency of haplotype *i* among contributors to sample *j* is

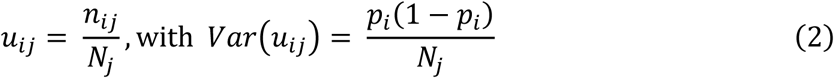

The vector of contributor haplotype frequencies, ***u_j_***=(*uij,…,ukj*), contains the signal of interest for estimating *N_j_* through its variance. However, ***u_j_*** is unobservable from eDNA samples.

Instead, we observe a downstream quantity: the number of sequencing reads for each haplotype, ***R_ij_***, and their relative read frequencies ***Rsub>ij*** =(*fij,…,fkj*). It is important to understand that these observed frequencies are not a direct reflection of ***u_j_***; they are influenced by a sequence of ecological and technical processes (Figure 2) that introduce additional noise and distort the original haplotype frequencies present among the contributors.

The first major source of variation arises from eDNA ecology (Barnes & Turner, 2016), where processes such as DNA production, degradation, and transport influence how much DNA each individual contributes to the local eDNA pool. This introduces variance, denoted here as *ε*_1_, between the true frequencies of haplotypes of contributors (*ui*) and the haplotype frequencies present in the water (*ri*; Figure 2b). Such variance depends on organism size, their distance to the sampling site, and hydrodynamic conditions (Yates et al., 2025). A second source of variance, denoted here as *ε*_2_, is introduced during sampling of the environment (Figure 2c). Only a fraction of the eDNA present in the water is collected, and the size and timing of this subsample influence the degree to which the sampled haplotype frequencies (*si*) frequencies present in the water. Larger sample volumes reduce this variance, but do not eliminate it. Subsequent laboratory steps introduce a third layer of variance, denoted here as *ε*_3_, affecting the transformation of sampled molecules into a DNA extract (*Ti*; Figure 2d). This includes variability from DNA extraction efficiency, pipetting error, PCR stochasticity, and differences in amplification efficiency among haplotypes. Finally, the sequencing process imposes multinomial sampling error denoted *ε*_4_ on the observed frequencies (*fi*; Figure 2e). This variance is inversely related to sequencing depth, and although it may be relatively small at high read counts, it still contributes to the overall divergence between observed and true haplotype frequencies.

Altogether, the variance in the final observed frequencies for each haplotype in each sample *fij* reflects the cumulative effect of all downstream processes following individual DNA shedding (Fig. 2f). Each step—DNA shedding, decay, capture, extraction, amplification, and sequencing—introduces additional stochasticity. Critically, these sources of noise are not necessarily biased with respect to the haplotype frequencies or identities present in the contributing individuals. If that assumption holds, then:

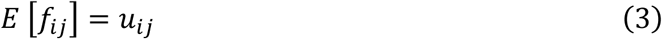

That is, the expected observed frequency still reflects the true underlying frequency among contributors, even if the observed values deviate due to added noise. In this view, the observed frequencies *f_ij_* are conditionally unbiased estimates of the unobserved *u_ij_*, but with added variance that is a function of the variance added in the processes highlighted above:

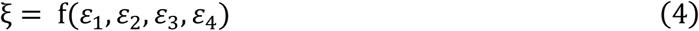

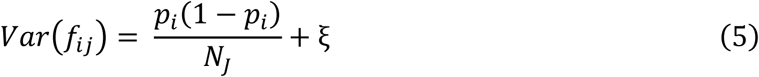

The additional variance ξ depends in part on the amount of DNA recovered in the sample, and may loosely correlate with the number of contributors, but it is not directly driven by it. As a result, it acts as an additive term that shifts the variance upward across all values of *N_j_*, and their impact is not negligible.

The key hypothesis is that ξ is small enough relative to the signal encoded in 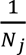 so that the original contributor number remains inferable. If the downstream noise dominates (e.g., due to low sampling volume or sequencing depth), then the observed haplotype frequencies become effectively random and independent of *N_j_*, and inference fails. In particular, when total variance at low contributor numbers becomes indistinguishable across samples, the method loses power to resolve small *N_j_* values. Thus, inference relies on the assumption that signal attenuation is incomplete and that the added noise does not overwhelm the relationship between variance and contributor number.

However, because biological replicate samples theoretically share the same local eDNA pool and observation processes, they offer a means of isolating the variance introduced by downstream processes. By quantifying between-sample variance across biological replicates (Figures 1c, 2c), or averaging across them, one can account for ξ. The residual variance—i.e., the difference between observed within-sample haplotype variance and population-level expectations—will then more closely reflect the biological signal of interest, allowing for improved estimation of the number of contributors.

### Deriving Population Haplotype Frequencies

Under the multinomial model described above (Eq. 1), the observed frequency of haplotype *i* in sample *j* a random variable with:

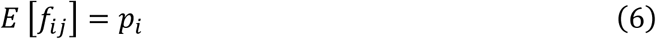

Because each observed *f_ij_* is an unbiased draw from a multinomial process centered on *p_i_*, averaging across multiple independent samples provides a natural estimator of the true population frequency:

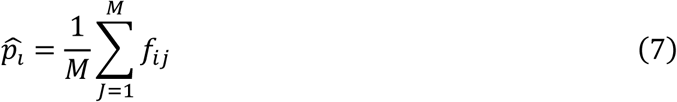

This estimator is unbiased, because *E* [*p̂_i_*] = *p_i_* . Thus, when multiple samples are considered, *p̂_i_* approximates *p_i_*. However, importantly, this averaging approach implicitly assumes that all samples contribute equally to the estimate of **π**—that is, it assumes approximate equivalence of Nj across samples. In theory, a more efficient estimator of **π** could be obtained by weighing samples based on their information content (e.g., read counts, total eDNA). Weighting by sequencing read depth may be tempting, but read counts are compositional, affected by variation in total DNA quantity, library normalization, and sequencing depth. As a result, read depth is not a reliable quantitative proxy for contributor number, particularly when species composition vary among samples or when multiple sequencing runs are used. Given these limitations, we default to using the equal-weight average across samples, which is unbiased and methodologically conservative. While this strategy does not minimize variance, it avoids introducing systematic distortions.

A practical concern arises when the same sample is used to both estimate π and infer its own number of contributors using the methods described below. In this case, the estimator is conditioned on a value it helps define, creating circularity. To avoid this, we apply a leave-one-out (LOO) approach: when estimating the number of contributors for a given sample j, we compute **π**^(−j)^, the estimate of population frequencies derived from all samples except j:

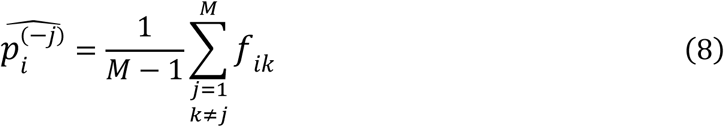

This ensures that the reference population frequencies used in estimating contributors to sample j are independent of the observed frequencies in that same sample. The estimator in Eq. 8 is the one used in downstream applications.

### Estimating Number of Contributors to Individual Samples

Once population haplotype frequencies are known, we may derive the number of individual contributors given the variability in observed frequencies. A statistically rigorous way to model the number of individual contributors in mixed eDNA samples would be to use a hierarchical model that explicitly accounts for variance added at each sampling stage highlighted in figure 2. For instance, biological replicates at the sampling stage (multiple bottle replicates) would allow estimation of variance arising from subsampling environmental DNA from the environment (Figure 2c). Then, technical replicates at the sequencing stage would enable quantification of variability introduced during laboratory handling and sequencing processes (Figure 2d–e). Once these sources of variance are estimated, the residual variance—the variance remaining unexplained—would more directly reflect the biological variance arising from the actual number of contributors to the eDNA sample (Figure 2a). Nonetheless, the variable contribution of individuals as in Figure 2b, arguably the highest source of variance, would remain.

In practice, the high level of replication required for a hierarchical approach is rare in most datasets, limiting its identifiability and applicability. Thus, while the model would be more accurate, it would not be useful in most instances. However, sampling, latent, and observation variances are largely independent of the true number of contributors (N), acting as noise that increases variance, thus artificially decreasing N estimates, but not erasing the relationship between estimated and true number of contributors. This allows us to simplify the model by ignoring these intermediate unobserved stages and their associated variances to directly link haplotype frequency variability in the eDNA metabarcoding data to biological variance in contributor numbers in the real world.

### Normal-Approximation Log-Likelihood Approach

Given Eq. 6, by the Central Limit Theorem (CLT), if the sample size is large enough, each observed frequency *f_ij_* can be approximated by a normal distribution with mean based on the population frequency *p_i_* and variance given by the multinomial process (Iliadis et al., 2012; Zhang et al., 2008):

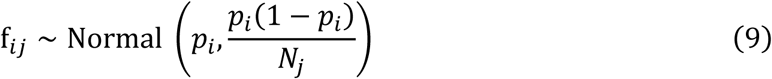

Under this assumption, then the probability density function (PDF) given by the normal distribution and the variance term is

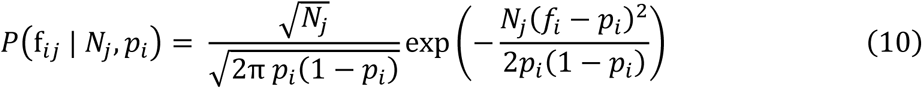

Then, naively assuming independence across k haplotypes, the log-likelihood is the sum of the log-PDFs:

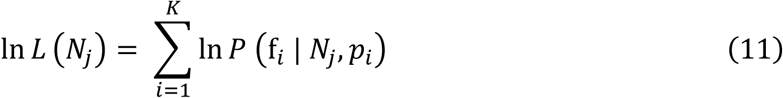

Substituting the PDF and simplifying, we get:

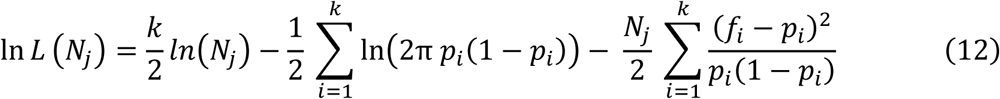

To maximize ln L(*N_j_*), we consider the negative likelihood, take the derivative of the N-dependent terms, and set them to zero. Fortunately, this leads to a closed form solution:

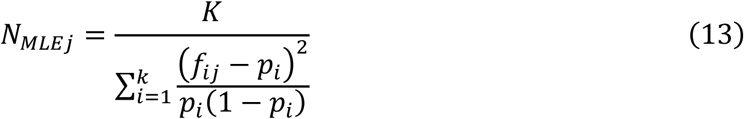

Where *N_MLRJ_* is a point estimate for the most likely number of contributors to an eDNA sample *j* given the total number of haplotypes *K*, a haplotyp’s frequency within the sample *f_ij_*, and the population frequency for that haplotype pi.

It is important to note that this method is a simplified approximation, because it ignores the covariance among haplotypes. This, however, brings a major advantage in allowing us to estimate *N_j_* from a single sample even with no replicates, and provides a likelihood framework that can be used to compare with other likelihood estimates. When multiple biological replicates are available, one may simply average their haplotype frequencies, and the result can be input in place of *f_ij_*. This is preferable, as the downstream sources of variance (Fig. 2c-e) are averaged upon, potentially reducing biases.

Finally, to assess uncertainty in N, we employ a profile likelihood approach (Murphy & and Van Der Vaart, 2000). Under standard regularity conditions, the likelihood ratio statistic −2[*lnL*(*N*) − ln *L*(*NN_MLE_*)] is asymptotically *X*^2^ distributed with 1 degree of freedom. We then define the confidence interval as the set of N values for which

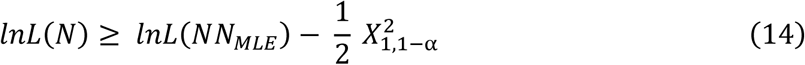

Where *X*_1,1−α_^2^ denotes the 100 (1 – α)% quantile of the chi-square distribution with 1 degree of freedom.

An alternative method based on a Method of Moments (MoM) approach, rather than a likelihood or normal approximation, is provided in the Supplement (Figure S3). However, because it consistently underperformed relative to the normal approximation across most scenarios, we do not present its results in the main text.

### Ploidy-based Minimum Number of Contributors

In addition to the likelihood approach, we also impose a minimum number of contributors to a sample, derived from the ploidy of the locus under analysis. Specifically, for haploid loci, we require that the estimated number of contributors to an eDNA sample must be at least equal to the number of observed haplotypes. For example, if four haplotypes are observed in the sample, regardless of their relative frequencies, the sample must have been derived from at least four distinct individuals. Similarly, for diploid loci, we define the minimum number of contributors as half the number of observed haplotypes, rounded up. Using the same example, a sample with four observed haplotypes could be explained by as few as two diploid individuals (e.g., two heterozygotes), but never fewer. Finally, in cases where ploidy cannot be inferred—such as with loci affected by heteroplasmy—this constraint can be disabled.

### Assumptions and Requirements

Beyond the requirement for known allele frequencies and the assumption that observation noise is not different for each haplotype, the method depends on other key assumptions to be laid out explicitly. First, it requires panmixia (i.e., no population structure within the study area) and that loci are in Hardy-Weinberg Equilibrium. These assumptions are crucial because processes such as local adaptation or population structure could cause certain alleles to be overrepresented in certain samples for reasons unrelated to sampling. To minimize this risk, the method should be applied to unstructured populations and focus on loci unlikely to be under strong selective pressure such as non-coding regions.

Second, it is assumed that Amplicon Sequence Variants (ASVs) observed in eDNA data correspond to real haplotypes in the population and are not false alleles (e.g., due to amplification or sequencing errors). These are potential concerns, as they have been observed in eDNA data (Elbrecht et al., 2018; Macé et al., 2022; Tsuji et al., 2020). Numerous approaches to correct for such errors are available, and should be applied to data before interpreting observed ASVs as haplotypes (Frøslev et al., 2017; Jeunen et al., 2024; Koseki et al., 2025).

Third, the method assumes that observed frequencies are centered around the frequencies of contributors (Eq. 3). Thus, it is also important that the sources of additional noise that are in control of the researcher be minimized. This can be done by maximizing sampled water volume, optimizing extraction efficiency, and maximizing sequencing read depth.

### Simulation

To validate our statistical methods, it was necessary to work with samples where the number of contributors was precisely known and controlled. To this end, we developed a series of three R functions (R core team, 2013) to simulate eDNA data and apply the method as a proof-of-concept. These functions allow us to generate synthetic datasets that replicate the key processes influencing eDNA sampling, including individual contributions, decay, allele frequencies, and sampling biases. The functions are potentially useful for other applications, and they are provided in the supplement and the github https://github.com/pedrobdfp/eDNA_haplo_frequencies.

First, the function generate_contributors randomly generates a number of individuals contributing to a local eDNA pool. For each individual k, it then assigns: (1) a haplotype identity via a multinomial draw from the user-defined population frequencies π; (2) a body size *Size_k_* drawn from a user-specified range; and (3) a distance *d*_*k*_ from the sample. Contributors may be generated as groups, in which case contributors share a single distance from the sample, or as ungrouped, in which case each contributor receives an independently sampled *d*_*k*_.

Then, the second function, generate_eDNA, takes in the output of the first function and simulates the process of eDNA generation and sampling from a local pool of contributors while keeping track of individual molecules. The function first calculates the eDNA shed for each contributor based on their body size and an allometric scaling exponent (default β = 0.75; Brown et al., 2004; Jo et al., 2024; Yates et al., 2025). Once the expected shedding rate is determined, the function reduces each contributor’s eDNA exponentially based on the distance *d*_*k*_ between that individual and the sampling location. Altogether, the process follows the equation described in Figure 2b. Following these steps, the function places molecules in a “local pool,” which keeps track of both the contributor ID and haplotype of each molecule.

The function then simulates the eDNA sampling process by drawing a fraction of the local pool into a “bottle replicate,” representing a subsample of the available eDNA in the environment. The number of molecules sampled is determined by a user-defined bottle_volume parameter, which specifies the proportion of the total local pool that is captured in each replicate. A random subset of molecules is selected without replacement, ensuring a realistic representation of stochastic sampling effects. For each replicate, the function summarizes the total number of eDNA molecules captured, the number of unique contributors present, and calculates haplotype-specific relative frequencies by counting how many molecules belong to each haplotype and normalizing by the total number of molecules in the sample. Thus, output consists of two tables: a summary table reporting total eDNA, number of contributors, distance metrics, and haplotype diversity for each sample and replicate; and a haplotype frequency table listing the relative abundance of each haplotype in each sample and replicate.

Finally, a third function, simulate_metabarcoding_data, takes the output of generate_eDNA and simulates high-throughput sequencing observations (i.e., metabarcoding reads) for each sample. This function draws a specified number of reads per sample using a multinomial distribution, where the probability of each haplotype is proportional to its relative abundance in the simulated eDNA pool. Crucially, because this simulation assumes uniform amplification efficiency across all haplotypes, the function does not have amplification steps. Additional stochasticity can be added via a normally distributed error term applied to the haplotype probabilities before sampling, mimicking noise introduced during library preparation. The function also supports multiple technical replicates per sample. All sources of variance including number of contributors, size, distance, decay rate, shedding rate, sequencing depth, and error can be specified, making this simulation framework ideal for testing the sensitivity and robustness of contributor estimation methods.

We conducted two types of simulations in this study: one under idealized “no-error” conditions and another using more biologically realistic parameters. For the no-error simulations (Figures 4, S4), we used deliberately simplified settings to isolate core model behavior. These included an arbitrarily high eDNA shedding rate (rate = 1000), no stochastic error (error = 0), high sequencing depth (100,000 reads), complete sampling of the collection volume (volume = 1), no DNA degradation (decay = 0), and equal DNA contribution from all individuals (no variation in contributor size and distance). For the realistic simulations (Figures 3, 5, S5), we introduced biologically plausible complexity. These runs used moderate process noise (error = 0.5), and sequencing depth typical for a single species in a multi-species metabarcoding dataset (1000 reads). We also introduced heterogeneity in individual contributors, with both contributor size and distance to the sampling point drawn from uniform distributions between 1 and 100. DNA degradation followed a realistic decay rate of 0.05 per unit of distance. Haplotype frequencies used across simulations varied, and are specified when relevant below.

**Figure 3:**
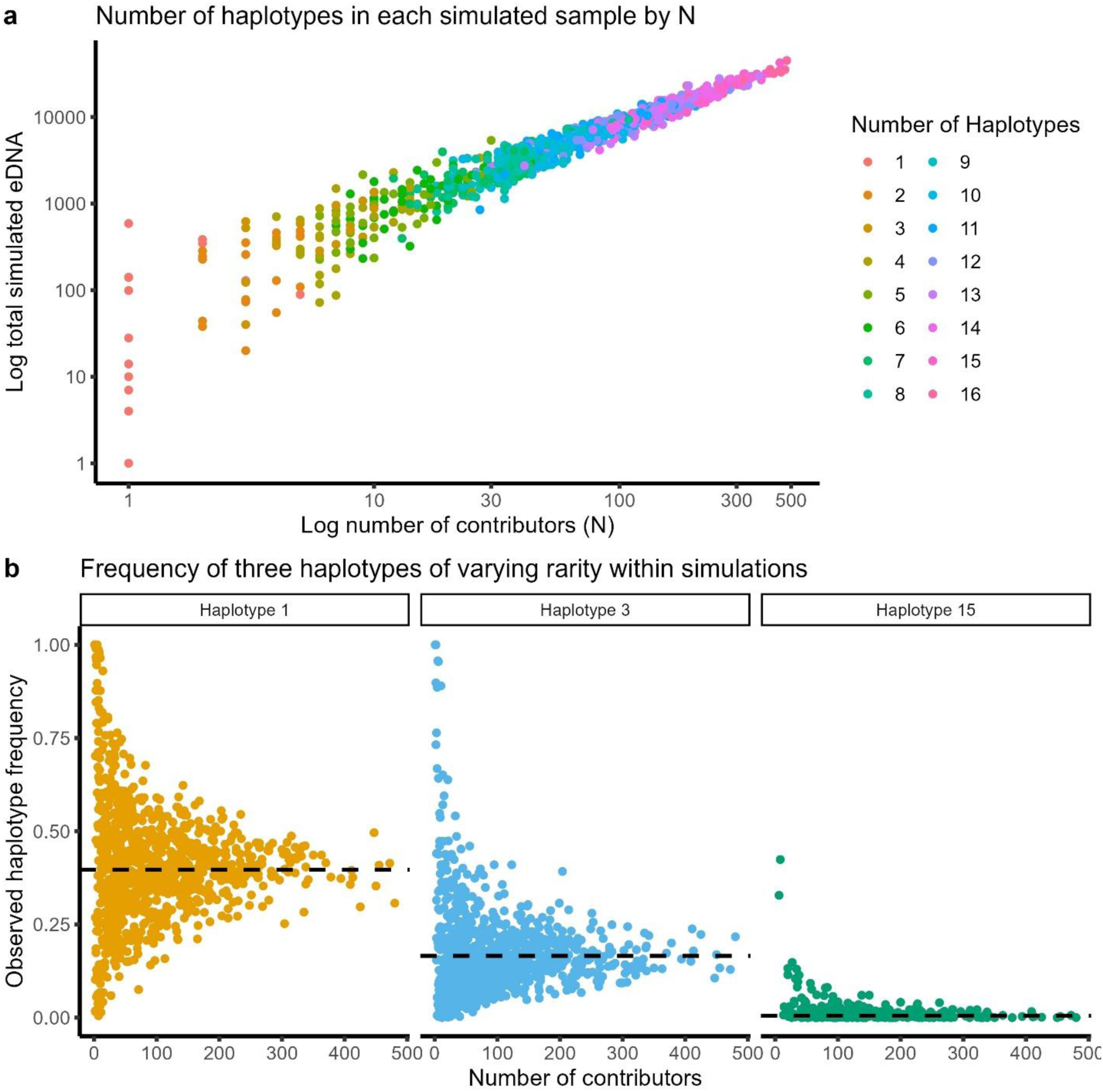
Haplotype patterns recovered from eDNA simulations. Simulations with a hypothetical variable marker with 16 alleles of varying frequency for ease of observation of patterns: π = (0.400, 0.165, 0.165, 0.050, 0.050, 0.050, 0.050, 0.010, 0.010, 0.010, 0.010, 0.010, 0.005, 0.005, 0.005, 0.005), 1000 samples with 1-500 contributors to each sample, that vary in size between 1-100. (a) Total simulated eDNA in each simulated sample by number of contributors with color showing number of haplotypes. (b) Relative frequency of three haplotypes of varying frequency within simulations. Haplotype 1 = 0.400; Haplotype 3 = 0.165; Haplotype 15 = 0.050. Dashed black line in each panel is the population frequency set for each haplotype. 1000 simulated samples plotted.

**Figure 4:**
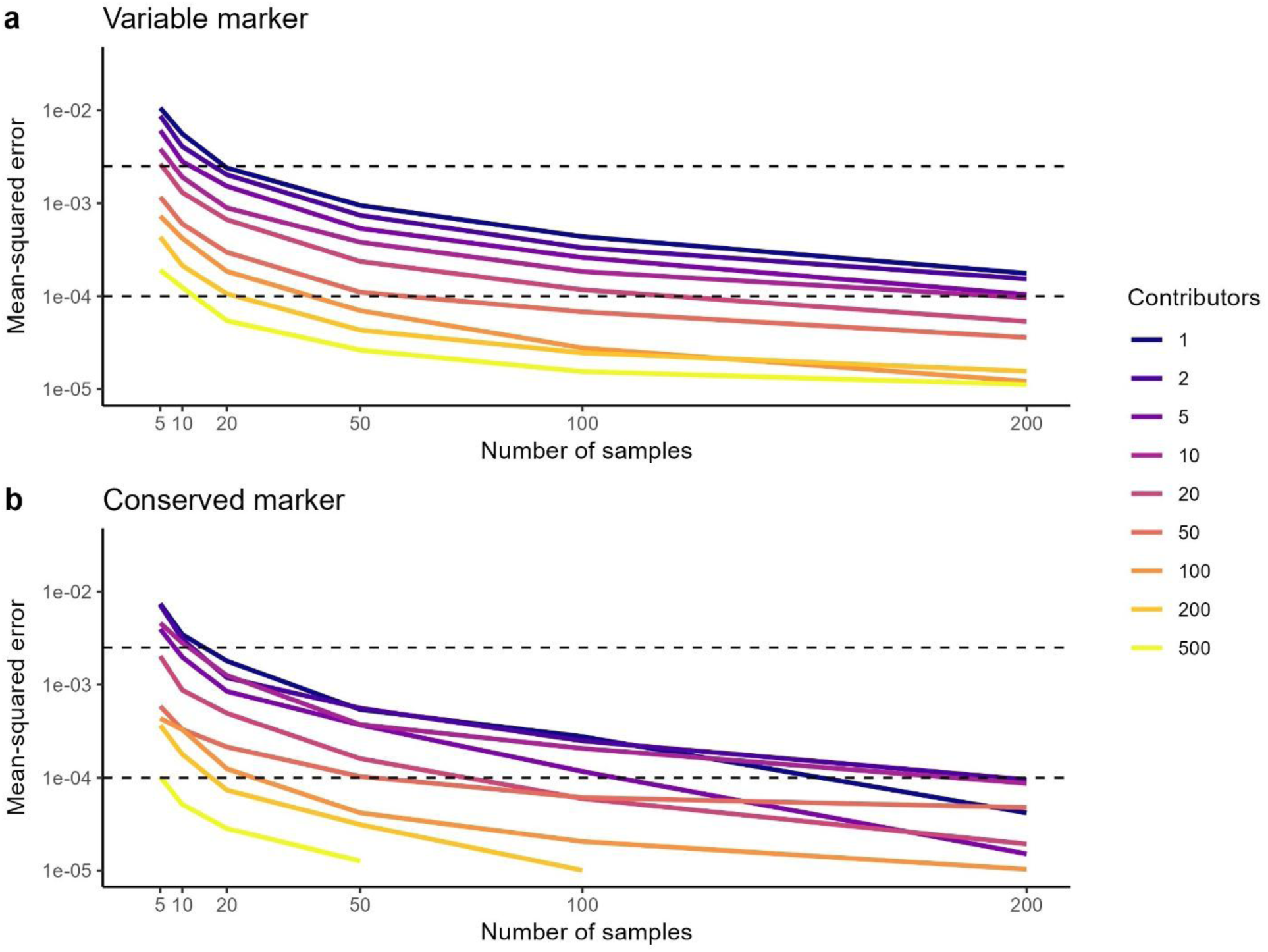
Error of population-frequency estimates as a function of sampling effort and mean number of DNA contributors. Lines show the mean-squared error (MSE) between the leave-one-out estimates of the population haplotype-frequency vector, Eq. 8, and the true vector (π) for an increasing number of independent eDNA samples. The horizontal dashed lines mark an MSE of 0.0025 (≈ 5 %) and 0.0001 (≈ 1%). (a) Variable-frequency mitochondrial marker. (b) Highly conserved mitochondrial marker. Colors differentiate scenarios with increasing average numbers of individual DNA contributors per sample (legend, right; darker blues = small local populations, lighter yellows = large). Simulations include the same level of variance as those in Fig. 6 (see below).

## RESULTS

### Environmental DNA Data Simulation

As expected, simulations of eDNA samples result in a variable number of haplotypes within samples, with more haplotypes observed as the number of contributors increases (Figure 3a). Furthermore, when we look at the variance in observed haplotype frequency as a function of the number of contributors (Figure 3b), the pattern depicted in Figure 2 emerges. We see the haplotype frequencies converging towards the overall population frequencies (dashed black lines), with stronger effect of sampling in samples with less contributors. This difference between observed frequency and population frequency is, in essence, what the proposed quantitative method uses to derive the most likely number of contributors.

### Estimating Population Frequencies from eDNA Samples

We assessed how accurately population-level haplotype frequencies (π) can be recovered from eDNA samples alone using the LOO averaging approach (Eq. 6). Simulations were performed under two scenarios: a hypothetical hypervariable marker comprising 17 haplotypes (π = 0.140, 0.120, 0.120, 0.100, 0.100, 0.080, 0.80 0.060, 0.060, 0.040, 0.040, v0.020, 0.020, 0.005, 0.005, 0.005, 0.005), and a conserved marker dominated by one out of six haplotypes (π = 0.86, 0.12, 0.005, 0.005, 0.005, 0.005). For each marker type, we generated synthetic eDNA datasets with varying average numbers of individual contributors per sample (1–200), who varied in their contribution to the sample by having variable size (1-100) and distance to source (1-100). We then evaluated how accurately population frequencies could be recovered as a function of the number of spatially replicated eDNA samples (Figure 4).

As expected, increasing the number of samples improved accuracy, and samples with more individual contributors yielded faster convergence toward true haplotype frequencies (Figure 4). For both marker types, frequency estimates stabilized rapidly: 20 samples were sufficient to recover population frequencies within a mean squared error (MSE) threshold equivalent to 5% deviation across haplotypes, regardless of the number of contributors or haplotype frequencies. When eDNA samples contained more than 50 contributors on average, population frequencies could be estimated within 1% error using as few as 50 samples. In all cases, eDNA-derived frequencies approximated true population frequencies more rapidly than tissue samples (Fig. 4, 1 contributor line). Nonetheless, for the conserved marker, luck in sampling rare alleles or not played an important role in the number of samples needed to approximate frequency (Fig. 4b)

### Accurate Prediction of the Number of Contributors of Simulated Data

To evaluate the best-case performance of our approaches, we first simulated eDNA data under ideal conditions (Figure 5): all individuals contributed equal amounts of DNA to the local eDNA pool, and no observational error was introduced (Figure 5b, d). Using haplotype frequencies from a hypothetical hypervariable marker with π = (0.140, 0.120, 0.120, 0.100, 0.100, 0.080, 0.080, 0.06, 0.06, 0.040, 0.040, 0.02, 0.02, 0.005, 0.005, 0.005, 0.005), the Normal Approximation Maximum Likelihood (MLE, Eq. 16) provided accurate estimates that closely matched the true number of contributors (Figure 5a). There was a strong correlation with true values (R = 0.849, p < 0.001), with 90% of estimates capturing the true value within the 95% confidence interval (coverage; Figure 5a).

**Figure 5:**
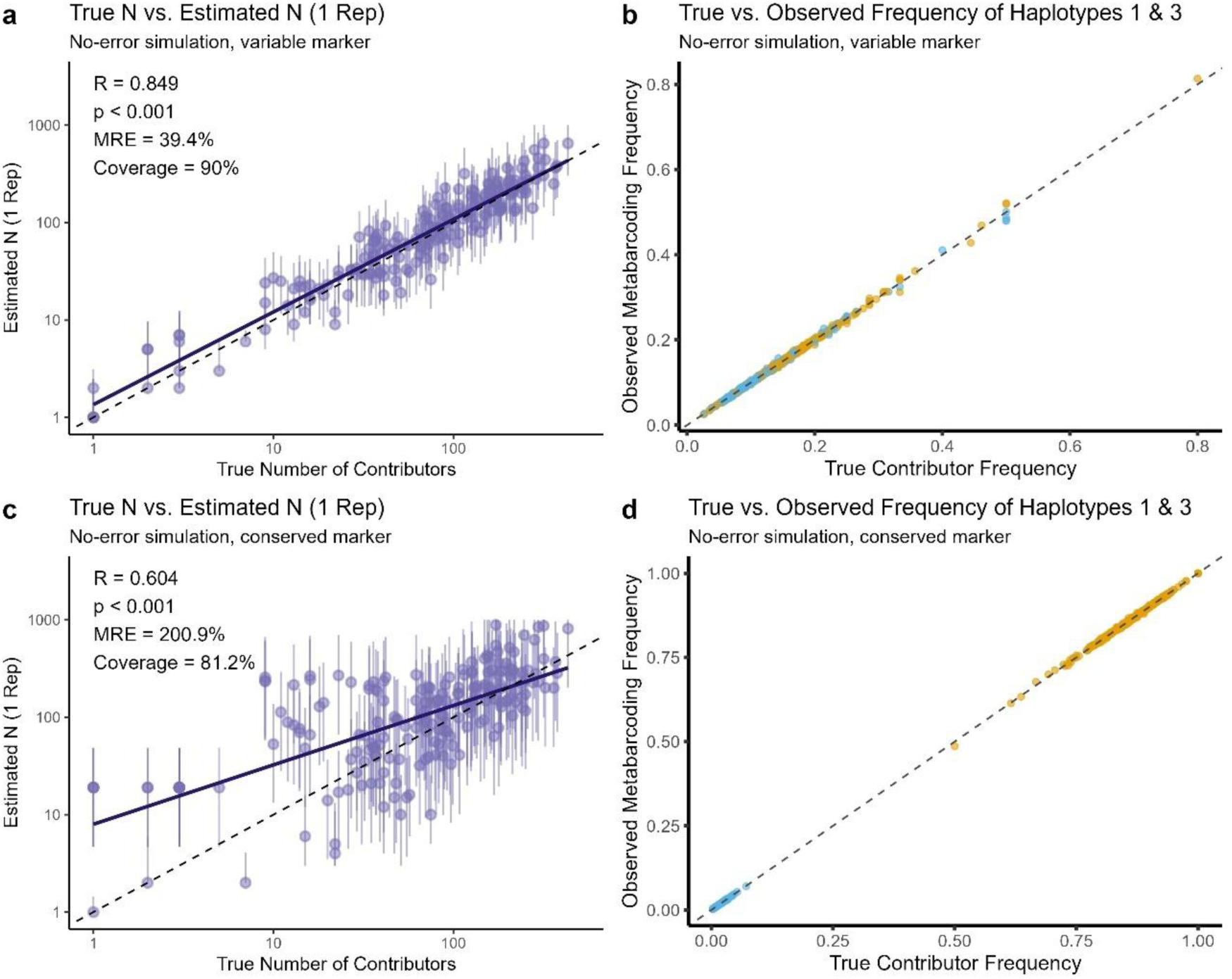
Precision of estimation method applied to simulated data with no added error. Panels (a–b) show results from a simulation using a hyper-variable marker. Panels (c-d) present a simulation using a conserved marker. Panels a and c show estimates derived using the Normal-Approximation Maximum Likelihood method (Eq. 13) compared to the true number of contributors to each simulated sample; panels (b, d) show the correlation between simulated contributors haplotype frequencies and the observed haplotype frequencies in the corresponding simulated metabarcoding observations, demonstrating that in these simulations, little error is added. Solid lines in all panels represent the linear regression fits, R is Pearson’s correlation coefficient, and dashed lines denote the 1:1 identity. Simulations included 5000 samples. While correlations are devised from all simulations, only a random subset of 200 are plotted here to avoid overplotting.

We then repeated the simulation using a marker with conserved haplotype characterized by a dominant allele and only 6 total haplotypes π = (0.86, 0.120, 0.005, 0.005, 0.005, 0.005). Under these conditions, the method produced strongly biased and less accurate estimates, which tended to be overestimated for lower numbers of contributors (Figure 5c). Correlation with the true number of contributors dropped to R = 0.604, MRE of 200.9%, and coverage to 81.2%. This is partially due to overestimation of samples containing only the most common haplotype. Because they have different numbers of contributors but only contain one and the same haplotype, they are all given identical, uniform estimations.

These results highlight that even under ideal conditions, the accuracy of contributor estimation is fundamentally tied to haplotype diversity at the locus. As haplotype distributions become increasingly skewed, particularly when one allele dominates, estimation error rises substantially (Figure S2).

### Additional Variance Effects on Estimations

We then repeated the simulations using both sets of allele frequencies, this time introducing high variability in individual eDNA contribution and some observation bias to assess how the methods perform under more realistic sources of error (Figure 6). Specifically, we incorporated heterogeneity in contributor size (ranging from 1 to 100, with larger individuals shedding more DNA following a 0.75 allometric scaling), much lower shedding rate to reduce overall DNA quantity (shedding rate = 5), variable distance to the sampling location (ranging from 1 to 100), with more distant individuals contributing less DNA due to decay (decay rate = 0.05 per unit of distance), eDNA pool subsampling (20%), lower sequencing depth (1000 reads) and some error (0.01) introduced by laboratory processes. This results in substantial differences between contributor haplotype frequencies and observed eDNA haplotype frequencies (Fig. 6c, f). We then repeated the estimator, using either one (Fig. 6a, d) or three (Fig 6b, e) biological replicates (independent bottle grabs simulated from the same local eDNA pool).

**Figure 6:**
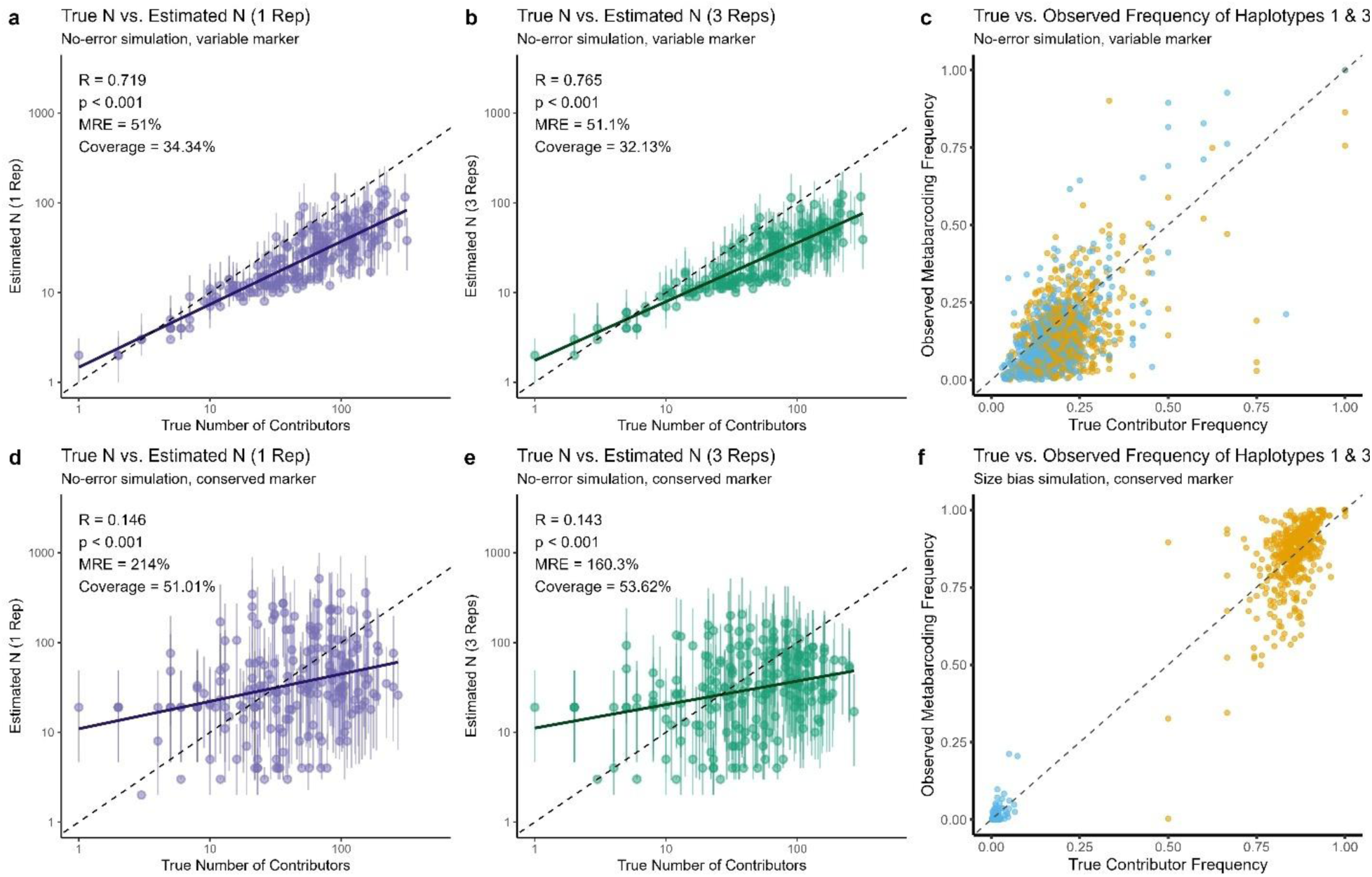
Precision of estimation methods applied to simulated data but with variable individual contribution. Panels (a–c) show results from a simulation using a hyper-variable marker (same frequencies as Fig. 5a-b) but with variable contribution from individuals including variable size (1-100) and distance from sample (1-100). (a) Estimation method applied to a single sample replicate with the hypervariable marker; (b) estimation method applied to three sample replicates with the hypervariable marker; (c) the correlation between simulated contributors haplotype frequencies and the observed haplotype frequencies in the corresponding simulated metabarcoding observations, demonstrating that in these simulations, each contributor provided variable amounts of eDNA to the sample resulting in differences between true and observed frequencies. Panels (d-f) present a simulation using a conserved marker (same frequencies as Fig. 4d-f) but otherwise identical simulation to previous panels. (d) Estimation method applied to a single sample replicate with the conserved marker; (e) estimation method applied to three sample replicates with the conserved marker; (f) same as panel c but with conserved marker. Solid lines in all panels represent the linear regression fits, R is Pearson’s correlation, and dashed lines denote the 1:1 identity. Simulations included 5000 samples. While correlations are devised from all simulations, only a random subset of 200 are plotted here to avoid overplotting.

As expected, the method underestimated the number of contributors under this scenario, especially in the samples with most contributors. This is due to error in frequency getting misinterpreted as a lower number of contributors. Nonetheless, estimates remained reasonable under the variable marker (Fig. 6a-c). When analyzing a single biological replicate (Fig. 6a), correlation with the true number of contributors dropped to R = 0.719 (a drop of – 0.130), MRE increased to 51% (a change of only +11.6%), but coverage dropped more significantly, with the correct answer recovered in only 34% of confidence intervals.

As discussed above, when observation noise is present, biological replicates can theoretically help remediate its effects, as the downstream noise (all except for eDNA ecology, Fig. 2c-e) can be averaged over. Accordingly, when considering three biological replicates (Fig. 6b), correlation is recovered slightly, but error as coverage remain not as good This is because the majority of differences between contributor haplotype frequencies and observed frequencies in the simulations were due to eDNA ecology (variable contribution from each individual), and not by laboratory and observation processes.

When using a combination of noise and a conserved marker (Figure 6d-f), the added noise worsened the estimations further, a result of removing the little signal present for these markers. The correlation between true and estimated numbers of contributors dropped substantially to only R =0.146, but remained significant (p <0.001); and coverage also dropped significantly to 51%. Much like the conserved marker, adding biological replicates had little effect in improving estimates (Fig. 5e).

For both sets of haplotype frequencies we observed a slope relative to the 1:1 line, where the number of contributors was slightly overestimated at low N. This is again caused by singletons of the most frequent haplotype.

## DISCUSSION

This study introduces a new framework for estimating the number of individual contributors to environmental DNA (eDNA) samples using within-sample haplotype frequencies and known or inferred population-level haplotype distributions. By leveraging the natural statistical convergence between observed and population haplotype frequencies, we show that it is possible to derive meaningful abundance estimates from eDNA metabarcoding data without directly relying on quantity of eDNA molecules. Simulations confirm that the method performs well under idealized conditions and remains informative even when incorporating biologically realistic sources of noise such as variation in contributor size, distance, and shedding rates. Together, these results suggest that haplotype frequency patterns offer a largely untapped source of quantitative information in eDNA data, bridging the gap between traditional presence/absence interpretation of metabarcoding and more species-specific quantitative approaches.

A key methodological advance introduced in our work, relative to existing approaches in the literature, is the use of the third tier of information: within-sample haplotype frequencies to estimate contributor abundance. Earlier models rely exclusively on the identity (Andres et al., 2021) or number (Ai et al., 2025) of haplotypes while ignoring the observed frequencies. Thus, our approach fully leverages the information present in the data, leading to more accurate and nuanced estimates. Notably, we do not explicitly model haplotype presence or combinatorial arrangements; instead, these effects are implicitly captured within the frequency patterns. This requires the assumption that observed frequencies are centered on the true frequencies of contributors. However, our simulations, which incorporate substantial and realistic levels of noise, show that the method is robust even when this assumption is moderately violated. It is possible, however, that integrating this method with previous methods might improve estimations.

We also show that the method is broadly applicable across loci, but performs substantially better when applied to hypervariable markers (Figs. 5, 6). This pattern mirrors that of previously proposed methods based on haplotype counts or identity alone (Andres et al., 2021; Yoshitake et al., 2019). The reliance on marker variability presents a challenge for applying the method to many existing eDNA datasets, as commonly used markers (e.g., 12S and COI) are relatively conserved. This tradeoff reflects a tradeoff within eDNA metabarcoding: to detect a broad range of taxa, primers must target conserved regions, which limits taxonomic resolution and the richness of quantitative signals. However, as sequencing technologies advance and longer reads can be reliably obtained from eDNA, more variable sites will be recoverable, increasing the power of haplotype-based inference.

In addition to estimating the number of contributors to individual eDNA samples, we also provide a framework for estimating population haplotype frequencies directly from eDNA data. While existing tissue-derived reference datasets could be used, estimating π from eDNA offers several methodological advantages. First, it ensures internal consistency: the same eDNA data used to estimate contributors also defines the reference frequencies. This alignment naturally captures any dataset-specific amplification or sequencing bias— biases that tissue-derived haplotype frequencies may miss, especially when primer mismatches or multiple sequencing runs are involved. Second, eDNA often recovers haplotypes that are rare or absent from tissue datasets (e.g., Parsons et al., 2025). When such haplotypes appear in subsequent samples, their lack of reference frequency renders them unusable unless population frequencies are inferred from the eDNA data itself. While no single sample is expected to capture complete haplotype diversity, pooled frequencies across spatially and temporally replicated samples can serve as a biologically and methodologically coherent proxy. This approach mirrors traditional population surveys, where reliability emerges from replication, not from any single observation.

### Future Steps and Validation

Here, we applied the method to simulated eDNA samples, but further method validation with field and real eDNA samples are necessary and encouraged prior to broad method application, as the level of “noise” added into real eDNA samples could be too high for this to be applicable in practice. The ideal dataset to validate the approach herein would include a population with known haplotype frequencies, a known or independently estimated number of contributing individuals, spatially distributed sampling, and eDNA sequencing with a hypervariable marker. To our knowledge, no publicly available dataset meets all of these criteria, and generating such a dataset is beyond the scope of this study. Most existing eDNA metabarcoding efforts rely on relatively conserved mitochondrial loci— such as 12S, COI, or CytB—chosen for their broad taxonomic coverage rather than their intraspecific resolution. While our method is theoretically compatible with these loci, our simulations demonstrate that its accuracy improves with more polymorphic markers (Figures 4–5), which provide finer resolution for distinguishing individual contributors. Thus, we judged it more appropriate to verify its effectiveness under simulated data only.

Future refinements could also improve abundance estimation by integrating haplotype-based inference with other independent data streams. One promising avenue is to combine this approach with total eDNA quantification (e.g., from qPCR or dPCR), which reflects bulk biomass but is subject to spatiotemporal biases (Ai et al., 2025). Because the two approaches capture distinct biological signals (number of individuals vs biomass) they can provide complementary and independent estimates of abundance (see Figure S4). In principle, a sample with high eDNA quantity but low haplotype diversity suggests many molecules from few individuals, whereas high values for both suggest contributions from many individuals. Additionally, running the method across multiple unlinked loci could further improve robustness, especially in diploid or recombining organisms. While we did not explore such integrative frameworks here, they represent a clear next step toward more accurate and biologically meaningful estimation of population abundance from eDNA.

### Method limitations

Our method relies fundamentally on the assumption that eDNA contributors represent independent random draws from an unstructured population. However, this assumption can be easily violated when natural populations exhibit familial clustering or complex population structure. For instance, it I common that invasive or expanding species will show spatial sorting in the edges of their distribution (Clarke et al., 2019; Comerford et al., 2023; Ochocki & Miller, 2017), which will break this assumption. Also, in species with strong social bonds or family groups, eDNA samples might capture genetic material from related individuals whose haplotypes are not independent. This is particularly problematic when analyzing mitochondrial markers, which follow strict maternal inheritance patterns. In such cases, our model would be unable to distinguish between different scenarios with the same genetic composition. For example, a single adult female versus a mother-calf pair sharing identical mitochondrial haplotypes, or a school of fish fry who are all siblings. Similarly, in populations with pronounced geographic structure, local samples may reflect haplotype frequencies that deviate systematically from the overall population frequencies (e.g. several Marine Mammals; Parsons et al., 2013, 2025), leading to biased contributor estimates. These biological realities highlight the importance of careful marker selection and thoughtful interpretation of results, particularly when applying this method to species with known social structures, philopatry, or during known reproduction events. Future refinements could incorporate prior knowledge of kinship patterns or spatial genetic structure to improve the accuracy of contributor estimates in such scenarios.

Additionally, as discussed in the Methods, a common limitation of approaches for estimating the number of contributors, including those presented here and those developed by Andres et al. (2021, 2023), is their reliance on population-level haplotype frequencies. In this study, we suggest approximating these frequencies using the distribution of haplotypes across multiple eDNA samples, treating the pooled frequencies as a proxy for population-level values. While we show this approach results in rapid approximation of true frequencies (Fig. 4), this must be done carefully. First, one must be aware of the potential inclusion of false or artifactual haplotypes caused by PCR or sequencing errors. To mitigate this risk when applying the methods herein to real datasets, it is important to use filtering methods that ensure observed sequence variants indeed relate to haplotypes. Even with stringent bioinformatic pipelines, false alleles may persist in eDNA data (Tsuji et al., 2020). Fortunately, tools like lulu (Frøslev et al., 2017) can collapse erroneous variants by clustering sequences based on similarity and co-occurrence, especially when calibrated with a mock community. More recently, dedicated R packages such as TombRaider (Jeunen et al., 2024) and gmmDenoise (Koseki et al., n.d.) have introduced statistical filtering tailored to eDNA data, which also remove spurious ASVs. Together, these methods make it practical to remove artifacts and recover reliable haplotypes from environmental samples. Second, as discussed above, deriving haplotype frequencies exclusively from eDNA demands a well-replicated sampling design that is spatially and/ or temporally distributed.

Finally, when it comes to interpreting the results, it is crucial to emphasize that the derived contributor numbers should not be interpreted as exact counts of individual organisms in an area. Rather, they represent an underrepresented estimation of contributors whose DNA was captured and retained through the sampling and sequencing process. This value relates only indirectly—and likely through a complex, nonlinear function—to true organismal abundance. For example, if two lakes are sampled for eDNA, and one lake yields an estimated 10 contributors and another 100, it is reasonable to infer that the latter harbors more individuals, but not necessarily exactly 100 individuals or even 10 times as many as the first. The method is best interpreted as providing a conservative lower bound on contributor number, excluding key real-world sources of variance such as differential shedding rates, environmental degradation, and sequencing bias.

## ACKNOWLEDGMENTS

Authors thank Amy Willis, Eric Ward, and Andrew Olaf Shelton for comments on statistical methods that resulted in several iterations and improvements. We also thank Kim Parsons, Elizabeth Brasseale, and other members of the Kelly lab and ‘Science Friday’ attendees for their helpful comments on early stages of the concept. This research was funded by the Office of Naval Research under Award Number (N00014-22-1-2719).

## DATA ACCESSIBILITY STATEMENT

All code necessary to replicate the simulations and a function to apply the likelihood method is made available in a quarto file in https://github.com/pedrobdfp/eDNA_haplo_frequencies, and as a supplement.

## BENEFIT-SHARING STATEMENT

This research contributes benefits by making all code publicly available through the databases described above.

## AUTHOR CONTRIBUTIONS

PBDFP conceived the study. All authors contributed to the development of methods and statistical approaches. PBDFP wrote the first draft of the manuscript, and all authors provided edits and approved the final version. EAA and RPK secured funding.

