## Supporting Information for "Estimating Organism Abundance Using Within-Sample Haplotype Frequencies of eDNA Metabarcoding Data"

### Table of Contents:


| | |
|---|---|
| **Figure S1** | Page 2 |
| **Figure S2** | Page 3 |
| **Method of Moments** | Page 4 |
| **Figure S3** | Page 5 |
| **Figure S4** | Page 6 |
| **References** | Page 7 |

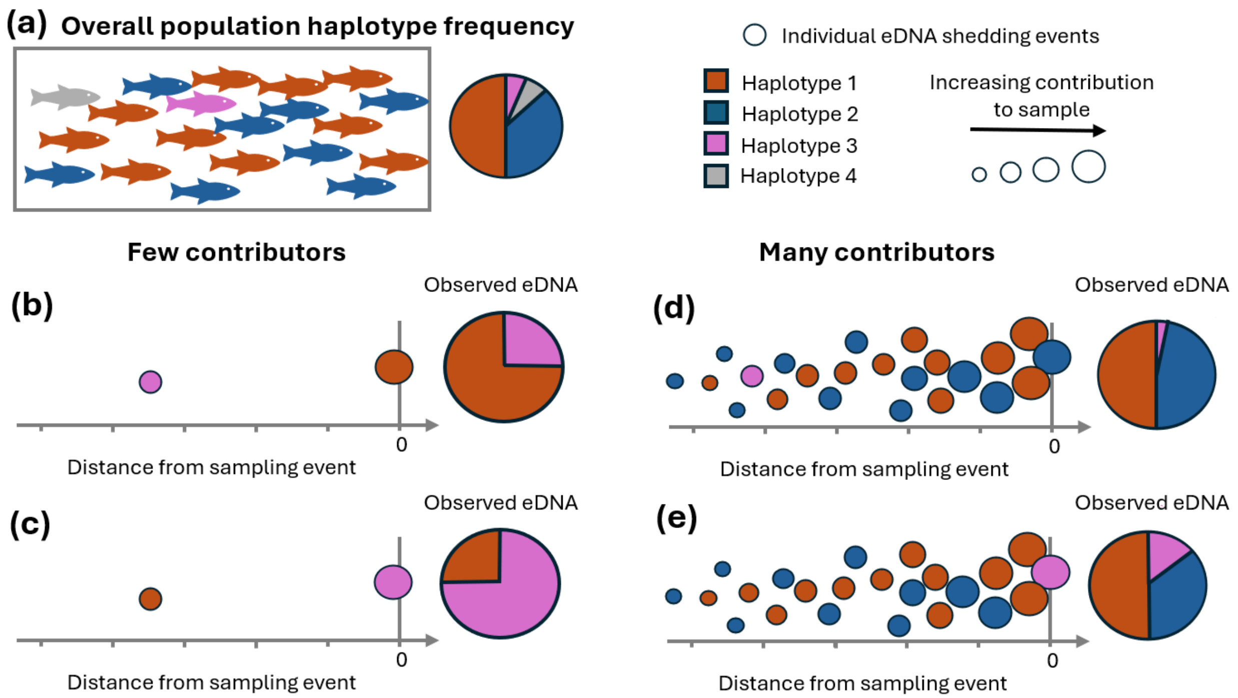


**Figure S1: Mechanisms driving the convergence of observed eDNA haplotype frequencies toward population frequencies.** (a) Population-wide haplotype frequencies: The pie chart on the right depicts the population haplotype frequencies, corresponding to the distribution of haplotypes (Haplotypes 1–4) represented by colored fish. (b-e) Individual eDNA shedding events: Circles represent individual eDNA shedding events, with circle size indicating their relative contribution to the sample, which is given by their spatiotemporal distance from the sampling location (distance = 0). (b, c) Small number of contributors: With less contributors, the observed haplotype frequencies are quite different from the population frequencies. In (b), Haplotype 1 (orange) dominates due to a nearby shedding event, causing observed haplotype frequencies to deviate significantly from population frequencies. Similarly, in (c), Haplotype 3 (pink) is overrepresented because of proximity to the sampling location. (d, e) Large number of contributors: With more contributors, eDNA from a greater diversity of individuals is captured, and observed haplotype frequencies increasingly align with population frequencies. The influence of a single contributor with Haplotype 3 (pink) becomes less pronounced, whether the individual is far from (d) or close to (e) the sampling location. Therefore, when fewer contributors are present (b, c), a single shedding event's position can have a disproportionate effect. In contrast, with many contributors (d, e), the aggregated contributions make the relative position of each individual event less impactful.

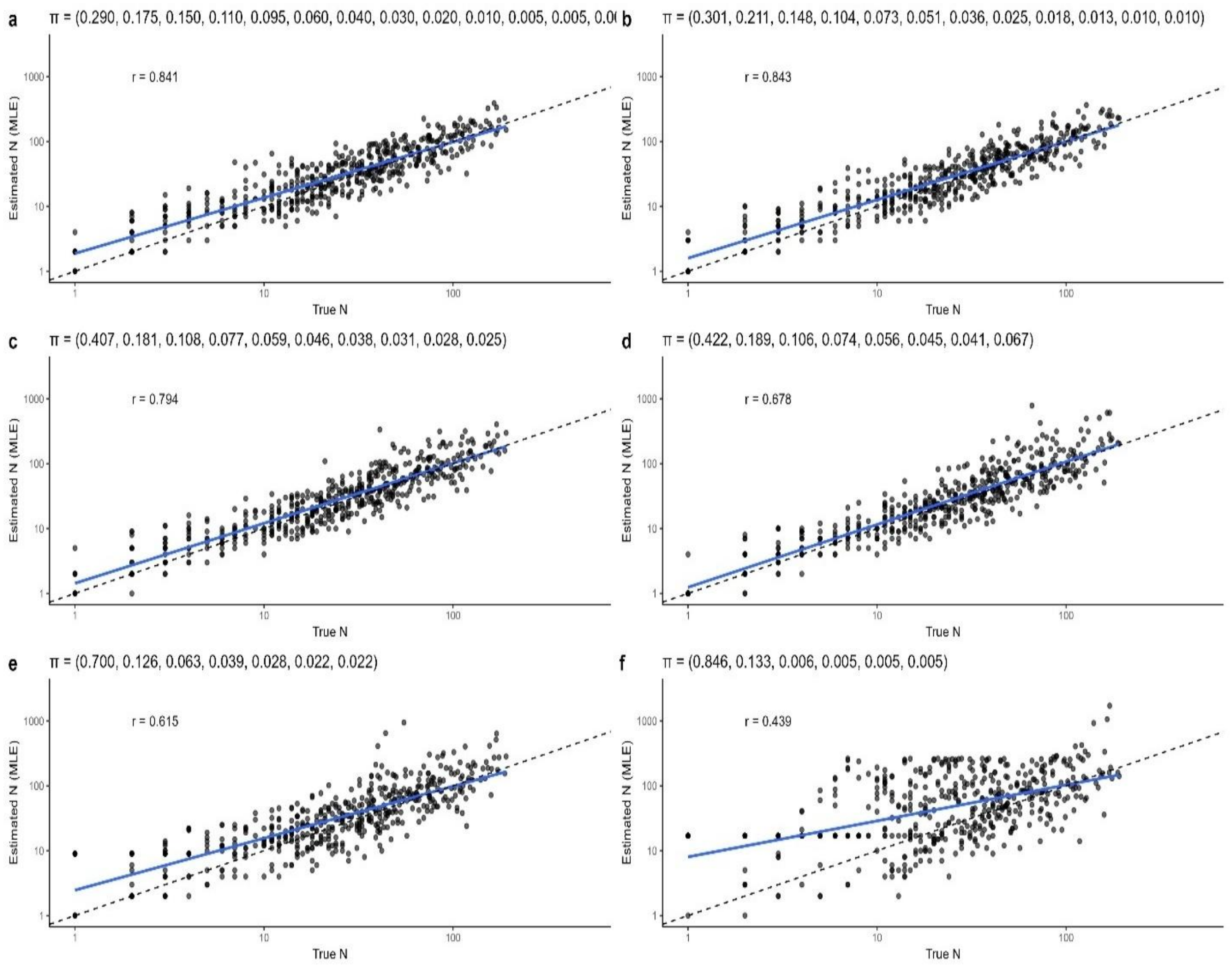


**Figure S2: The effect of various haplotype frequencies (π) on effectiveness of estimations.** π above each pair of panels show the haplotype frequencies in the simulations. R is Pearson's correlation, dashed line is the 1:1 line. Simulations performed with no error to determine maximum effectiveness of the method

### Method of Moments Approach

An alternative to the normal approximation maximum likelihood approach presented in the main text is the method-of-moments approach. It is mathematically equivalent, as it stems from the same principle and the final derived equation is very similar. Nonetheless, the method does not provide a likelihood distribution, making it challenging to combine multiple loci. Additionally, preliminary tests showed that the Normal MLE approach performed better under most scenarios (Figure S3). Hence, it was presented in the main text in favor of this method. Nonetheless, we chose to present the method of moments solution here as well, as it may be useful for integration into other non-likelihood frameworks.

Under the multinomial model (Eq. 1-4 in the main text), the expected sum of squared deviations between the sampled frequencies $f$ and the population frequencies π, that is, the difference between observed and population haplotype frequencies, is given by equation 7:

$$E\left[\sum_{i=1}^{K}(f_i - p_i)^2\right] = \frac{1 - \sum_{i=1}^{K} {p_i}^2}{N} \qquad (S1)$$

In essence, it states that as N increases, the observed frequencies $f$ become closer to π, and the total deviation shrinks proportionally to 1/N. With known population frequencies, the square deviations of frequencies can be derived from samples:

$$S_{OBS} = \sum_{i=1}^{K}(f_i - p_i)^2 \qquad (S2)$$

Then, under the method of moments (Luikart et al., 1999; Osękowski, 2017; Waples, 1989), we equate the observed value $S_{OBS}$ (Eq. S2) to its theoretical expectation that is a known function of the unknown parameter (Eq. S1) Therefore:

$$\sum_{i=1}^{K}(f_i - p_i)^2 = \frac{1 - \sum_{i=1}^{K} {p_i}^2}{N} \qquad (S3)$$

Solving for N, we obtain

$$N_{MoM} = \frac{1 - \sum_{i=1}^{K} {p_i}^2}{\sum_{i=1}^{K}(f_i - p_i)^2} \qquad (S4)$$

Solving this equation with the observed frequencies of all haplotypes in the sample will provide a point estimate for the expected number of contributors to the eDNA sample, given the observed frequency variance. To provide confidence intervals, we can approximate the uncertainty in N via the delta method (Dorfman, 1938). In our implementation, the standard error (SE) of N is approximated as

$$SE(N_{ij}) \approx \sqrt{\frac{2N^2}{n}} \tag{S5}$$

Where n is the number of haplotypes with non-zero observed frequency. With this standard error, we construct a 100(1 – α) % confidence interval for N as

$$N_{ij} \pm z_{1-\alpha/2} \times SE(N_{ij}) \tag{S6}$$

Where z is the z score from the normal distribution.

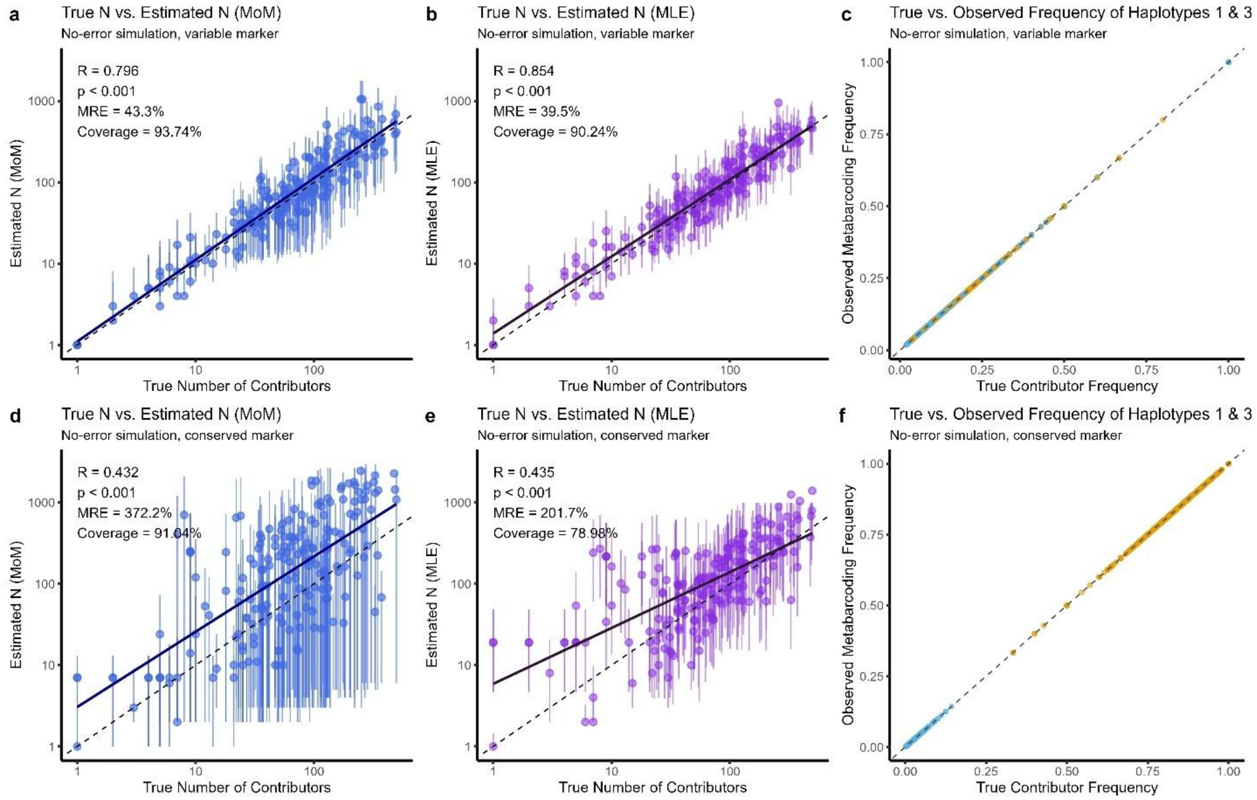


**Figure S3: Comparison Between Normal Approximation MLE and Method of Moments (MoM).** Panels (a–c) show results from a simulation using a hyper-variable marker. Panels (d-f) present a simulation using a conserved marker. In panels (a, d) estimates were derived using the Method of Moments (MoM, Equation S4); in panels (b, e) estimates were derived using the Normal-Approximation Maximum Likelihood method (main text); panels (c, f) show the correlation between simulated contributors haplotype frequencies and the observed haplotype frequencies in the corresponding simulated metabarcoding observations, demonstrating that in these simulations, little error is added. Solid lines in all panels represent the linear regression fits, R is Pearson's correlation, and dashed lines denote the 1:1 identity. Simulations included 5000 samples. While correlations are devised from all simulations, only a random subset of 200 are plotted here to avoid overplotting.

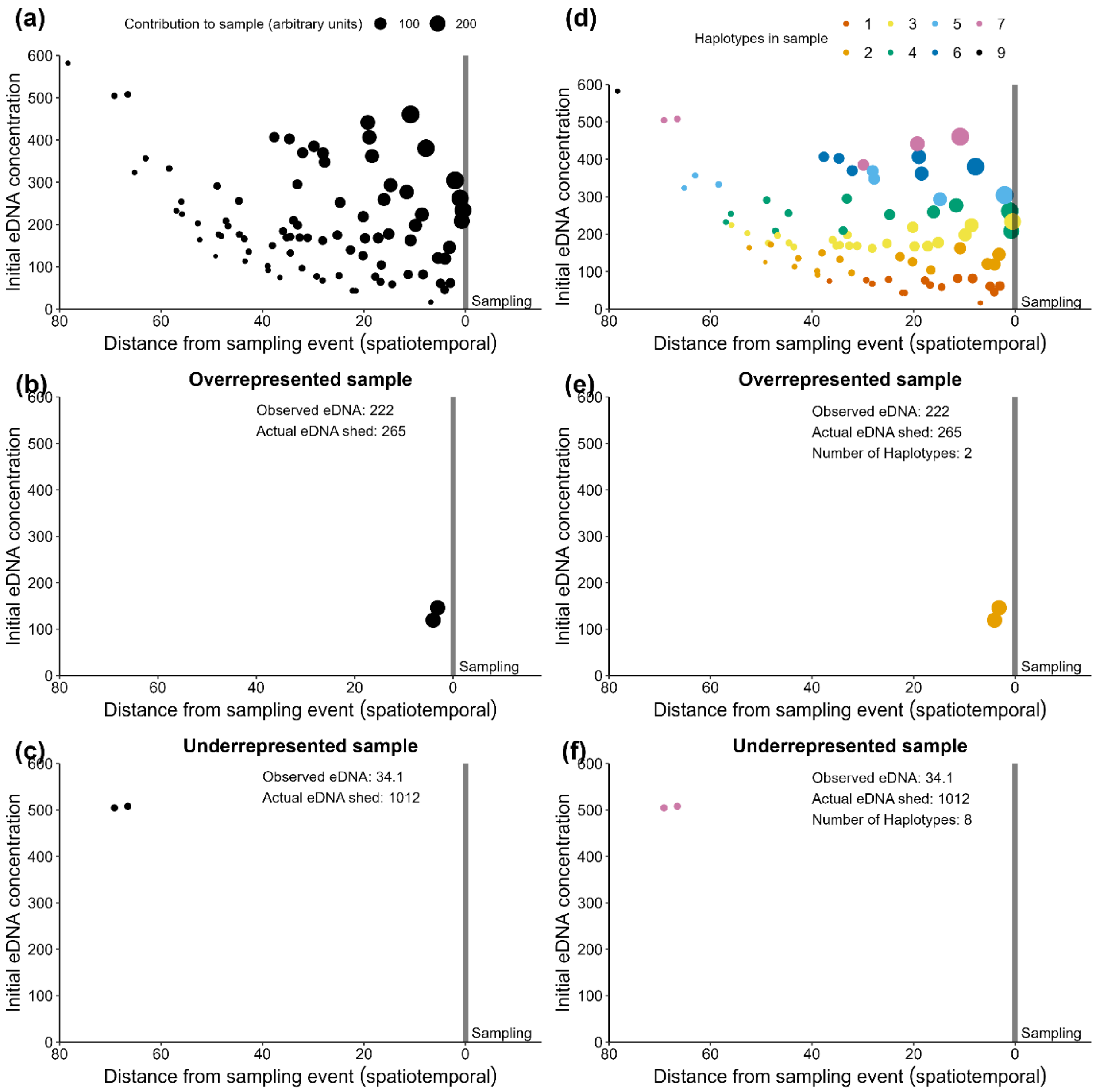


**Figure S4: Simulation figure illustrating how haplotype number and eDNA concentration can provide conflicting but complementary information.** In panel (a), each black dot represents an eDNA shedding event, varying in initial concentration and occurring at different distances—either spatially or temporally—from the sampling point. Consequently, when a sample is taken, these events contribute unevenly to the eDNA sample, leading to biased estimates that tend to favor the most recent or nearby shedding events. For instance, in panel (b), shedding events occur close to the sampling point, resulting in an overestimation of local target abundance if only DNA concentration is considered. Conversely, in panel (c), the shedding event occurs farther from the sampling point, leading to an underestimation of abundance if only concentration is considered. Therefore, if this bias can be corrected, then theoretically we would improve abundance estimates from eDNA data. As the number of individuals contributing to a sample increases, so does the haplotype diversity, as shown in panel (d). Therefore, in a scenario like panel (e), where observed eDNA concentration is high but haplotype diversity is low, we may infer that the sample is overrepresented due to shedding events occurring close to the sampling point. Conversely, in panel (f), where eDNA concentration is low but haplotype diversity is high, it indicates that the sample is underrepresented due to shedding events occurring farther from the sampling point.
